# Corncob Bedding Negatively Impacts Breeding Performance and Sexual Development in Mice

**DOI:** 10.64898/2026.01.22.701214

**Authors:** Victor Lujan, Anna S. Ratuski, Kyna A. Byrd, Kendall M. Coden, David E. Bentzel, Joseph P. Garner

## Abstract

Corncob bedding is commonly used for housing rodents in research, but previous work has linked corncob to altered reproductive behavior, disrupted estrous cycling, aggression, and welfare impacts across non-murine rodents. Furthermore, corncob is used as a licensed commercial rodenticide. Corncob contains endocrine-disrupting compounds (EDCs) that interfere with aromatase activity and estrogen signaling, processes critical for normal sexual behavior and development, yet effects on reproductive outcomes in mice remain unexplored. We conducted two experiments to test whether corncob bedding influences breeding performance and male sexual development. In Experiment 1, we analyzed breeding records to compare breeding performance of NSG mice housed on corncob *versus* cellulose bedding across two 3-month phases (N = 488 litters). Pairs housed on corncob produced significantly fewer pups than pairs housed on cellulose. To understand this effect, in Experiment 2, hormonal and morphological effects of corncob were assessed in male mice from four genetic backgrounds (C57BL/6, BALB/c, FVB, and CD1; N = 32 cages). Mice were bred and born on aspen or corncob, with half switched at weaning and half unchanged. Corncob produced timing-dependent effects in male reproductive physiology and development. Early-life corncob exposure altered baculum morphology and reduced testosterone, estradiol, and anogenital distance. In contrast, post-weaning corncob exposure resulted in hyper-masculinization, indicated by increased anogenital distance. Alongside prior evidence that corncob contains EDCs, our results raise serious concerns about its suitability as bedding in animal research. Continued use of corncob introduces uncontrolled variation that compromises animal welfare, reproduction, experimental validity, and reproducibility.

## Introduction

Corncob is widely utilized as a bedding material for rodents in research facilities throughout the United States (Villalon Landeros et al., 2012) due to its perceived benefits (e.g., cost, convenience of use in automated bedding dispensers). However, it has been shown in several rodent species that corncob negatively impacts animal welfare, reproductive behavior, and pathophysiology. For example, corncob bedding is associated with signs of stress such as increased mouse aggression and barbering, particularly in individually ventilated cages (IVCs) (Theil et al., 2020; Ratuski et al., 2025), elevated blood glucose in mice (Kondo et al., 2022), and reduced slow-wave sleep in rats (Leys et al., 2012). When given a choice, rodents prefer softer bedding alternatives over corncob, such as aspen wood shavings or paper-based options (Mulder, 1975; Krohn & Hansen, 2008). Corncob-derived cellulose has also been evaluated and regulated for use as an active substance in rodenticide (Health Canada PMRA, 2025), raising toxicological concerns. In addition to these welfare impacts, there is evidence that corncob exposure can lead to hormonal disruption with important impacts on endocrine function and sexual behavior shown in rats (Mani et al., 2005; Markaverich et al., 2005). Given that mice are the most used animals in preclinical research (European Commission, 2024; UK Home Office, 2023; Canadian Council on Animal Care, 2023), the lack of evidence regarding the effects of corncob bedding in mice represents a critical gap in the literature and positions corncob as an uncontrolled variable in mouse experiments and breeding operations.

Corncob contains endocrine-disrupting compounds (EDCs), including tetrahydrofuran-diols (THF-diols) and leukotoxin-diols (LTX-diols), which interfere with aromatase activity, effectively disrupting the conversion of testosterone into estradiol (Markaverich et al., 2005; Villalon Landeros et al., 2012). Exposure to LTX-diols is linked with altered estrous cycling in female rats, thereby impairing the conditions necessary for successful mating, ovulation, and conception (Markaverich et al., 2005). In male rats, exposure to THF-diols significantly reduces mounting, intromission frequency, and grooming behaviors (Mani et al., 2005). THF-diols also stimulate MCF-7 breast cancer cell proliferation in BALB/c mice, highlighting their broader pathophysiological impacts (Markaverich et al., 2002). Aromatase and estrogen play a critical role in shaping reproductive behavior and brain development, particularly through the actions of estrogen receptor alpha (ESR1). California mice (*Peromyscus californicus*) housed on corncob have fewer ESR1-positive cells in their brains, demonstrating a link between corncob exposure and altered neuroendocrine function (Villalon Landeros et al., 2012).

Sexual differentiation of genitalia is tightly regulated by hormonal signaling, primarily through both testosterone and estrogen—with estrogen produced from testosterone via the aromatase enzyme (Baskin et al., 2021). Disruption of aromatase activity during critical early developmental stages interferes with this hormone balance during a period of heightened endocrine sensitivity. At low doses, EDCs can alter gene expression and hormone signaling, leading to lasting developmental changes not seen in higher, toxic doses; these effects have been observed across laboratory animal models, particularly rodents and amphibians, and are supported by epidemiological evidence in humans (Ryan & Vandenbergh, 2002). As such, aromatase disruption during development is also associated with altered reproductive morphology, including hypospadias (a condition in which the urethral opening is abnormally located on the underside of the penis) and reduced anogenital distance (AGD) in humans, mice, and rats (Baskin et al., 2021; Schwartz et al., 2019). Anogenital distance is determined during a narrow prenatal window and is a well-established biomarker for endocrine disruption *in utero* (Schwartz et al., 2019).

The baculum, or “penis bone”, is also highly sensitive to endocrine signaling during development (Ghione et al., 2025). Androgen receptor activation in penile chondrocytes is essential for normal baculum formation: conditional androgen receptor knockout mice demonstrate underdeveloped bacula, reduced bone volume, and failed copulation, despite normal external genitalia and behavior (Ghione et al., 2025). In addition, variation in baculum morphology is associated with male reproductive success under competitive mating conditions (Stockley et al., 2013). Thus, AGD and baculum morphology are potential measures of the developmental impact of EDCs, including disrupted estrogen signaling related to corncob bedding exposure.

We conducted two experiments to test the effects of corncob bedding on reproductive development and performance in mice. Experiment 1 tested whether corncob bedding influences reproductive success in a large breeding colony of NSG mice. Breeding records were compared between mice switched to corncob bedding or those remaining on virgin paper cellulose bedding for 13 weeks, and again during a second 13-week phase when all mice were moved back to cellulose bedding. We hypothesized that mice housed on corncob bedding would exhibit reduced breeding performance.

Experiment 2 tested whether exposure to corncob bedding alters male reproductive physiology and morphology, and employed a 2×2×4 factorial design to test whether cage type (IVC vs static) and bedding type (corncob vs aspen) influence male reproductive development in four mouse strains. We measured blood testosterone and estradiol, along with AGD and baculum morphology. We hypothesized that mice housed on corncob bedding would show elevated testosterone, reduced estradiol, shorter AGD, and altered baculum morphology compared to mice housed on aspen bedding. Given organizational *versus* activational effects of steroid hormones, we anticipated that these effects might differ on the basis or pre-weaning and post-weaning exposure. Furthermore, given previous findings, we expected these effects to be amplified by IVC housing (Theil et al., 2020; Ratuski et al., 2025).

## Results

### Experiment 1

In Phase 1, there was a significant main effect of litter event (F_1, 322_ = 56.33, p < 0.0001), indicating that litter size differed between birth and weaning. There was also a significant main effect of bedding treatment (F_1, 322_ = 43.45, p < 0.0001), with mice housed on cellulose producing more pups than those housed on corncob bedding. The interaction between litter event (birth or weaning) and treatment was not statistically significant (F_1, 322_ = 0.17, p = 0.6770), indicating that the difference in pup loss from birth to weaning was similar in both treatment groups. Least Squares Means estimates (± SE) showed average pup counts of 7.12 (± 0.07) for litters born on cellulose, compared to 6.67 (± 0.06) for litters born on corncob. At weaning, there were an average of 6.61 (± 0.07) pups in litters on cellulose versus 6.21 (± 0.06) for litters weaned on corncob. A Least Significant Number power analysis revealed that a statistically significant interaction between litter event and treatment would have required 14,319 litters, suggesting that any true difference in pup mortality between treatments is likely minimal or biologically irrelevant.

In Phase 2, there was a significant main effect of litter event (F_1,162_ = 28.67, p < 0.0001) and breeding pair type (F_1,162_ = 60.52, p < 0.0001) on the number of pups per litter. On average, mice that remained on cellulose bedding throughout the study (i.e., cellulose-cellulose) had fewer pups than those that were exposed to corncob in Phase 1 but returned to cellulose in Phase 2 (i.e., corncob-cellulose). The interaction between litter event and breeding pair type was not statistically significant (F_1,162_ = 0.19, p = 0.6642), indicating that again, pup losses from birth to weaning were similar regardless of a breeding pair’s bedding history. After adjusting for seasonal or age-related decline observed between Phase 1 and Phase 2, litters from cellulose-cellulose pairs averaged 7.12 (± 0.09) pups at birth and 6.61 (± 0.09) at weaning, while corncob-cellulose litters rebounded to exceed the adjusted baseline, averaging 7.79 (± 0.07) pups at birth and 7.21 (± 0.07) at weaning. A Least Significant Number power analysis indicated that a statistically significant interaction between litter event and breeding pair type would have required a sample of 6,665 litters, indicating that differences in pup mortality between groups were again negligible.

### Experiment 2

There was a significant main effect of pre-weaning bedding on estradiol levels (F_1,128_ = 4.02, p = 0.0472), with males born on aspen exhibiting higher estradiol levels than those born on corncob. Post-weaning bedding alone did not have a significant main effect on estradiol (F_1,128_ = 0.87, p = 0.3520). There was a significant interaction between pre-weaning and post-weaning bedding (F_1,128_ = 14.23, p = 0.0002), indicating that the effect of post-weaning bedding on estradiol levels depended on the bedding that males were exposed to early in life. Among males born on aspen, those weaned onto aspen showed significantly higher estradiol levels than those weaned onto corncob (F_1,128_ = 11.08, p = 0.0011). Among males born on corncob, there was no significant effect of post-weaning bedding. Furthermore, males born and weaned on aspen had higher estradiol levels than males born on corncob and weaned onto aspen. In other words, any exposure to corncob reduced estradiol. Increases in estradiol in animals housed on aspen were greatest in IVC cages (interaction F_1,128_ = 4.36, p = 0.0388)

There was a significant main effect of pre-weaning bedding on testosterone levels (F_1,128_ = 10.99, p = 0.0012), with males born on aspen exhibiting higher testosterone than those born on corncob. Post-weaning bedding alone did not significantly influence testosterone levels (F_1,128_ = 0.40, p = 0.5303). There was a significant interaction between pre-weaning and post-weaning bedding (F_1,128_ = 8.01, p = 0.0054), demonstrating that the effect of post-weaning bedding depended on the bedding that males were exposed to early in life. Among males weaned onto aspen, those born on aspen showed significantly higher testosterone levels than those born on corncob (F_1,128_ = 18.88, p < 0.0001). Among males born on corncob, those weaned onto corncob showed significantly higher testosterone levels than those weaned onto aspen (F_1,128_ = 5.98, p = 0.0158). In contrast, post-weaning bedding did not significantly affect testosterone levels among males born on aspen (F_1,128_ = 2.42, p = 0.1223). There was no effect of cage ventilation on testosterone levels (interaction F_1,128_ = 0.07, p = 0.7869).

There was a significant main effect of pre-weaning bedding (F_1,127_ = 19.04, p < 0.0001) on AGD, with males born on aspen exhibiting a larger AGD (1.2455 ± 0.0026 mm) than those born on corncob (1.2296 ± 0.0026 mm). There was a significant main effect of post-weaning bedding (F_1,127_ = 21.74, p < 0.0001), with males weaned onto corncob having a larger AGD (1.2464 ± 0.0026 mm) than males weaned onto aspen (1.2273 ± 0.0026 mm). There was no significant interaction between pre-weaning and post-weaning bedding on AGD (F_1,127_ = 0.35, p = 0.5533). Animals housed on aspen did not differ between IVCs and static cages. Animals housed on corncob had significantly higher AGD in IVCs *versus* static cages (interaction F_1,127_ = 6.66, p = 0.0110).

Pre-weaning bedding affected baculum morphology (F₅,₇₅₄ = 5.32, p < 0.0001). There was a significant difference in the angle of curvature for the shaft (F₁,₉₀₀ = 17.94, p < 0.0001), with males born on corncob having a larger normalized angle of curvature than males born on aspen. Similarly, there was a significant difference in the normalized angle of curvature for the bulb (F₁,₉₀₀ = 5.42, p = 0.0201) with corncob males having a larger angle than aspen males. For the normalized extension of the bulb along the shaft, corncob males had a shorter extension than aspen males (F₁,₈₉₉ = 5.76, p = 0.0166). There were no significant differences in total baculum length (F₁,₈₉₉ = 0.45, p = 0.5023), bulb width (F₁,₈₉₉ = 2.70, p = 0.1006), or shaft width (F₁,₈₉₉ = 2.82, p = 0.0932) between bedding treatments.

## Discussion

The results of this study demonstrate that corncob bedding negatively impacts reproductive success, reproductive physiology, and sexual development in mice. We first show that corncob bedding reduces breeding performance in NSG mice during active exposure. Second, we show that corncob induces time-dependent changes in male reproductive morphology and hormone levels across four commonly used strains. In Experiment 1, exposure to corncob bedding during breeding resulted in fewer pups born and weaned, effects that were reversible upon return to cellulose bedding. In Experiment 2, exposure to corncob bedding during critical developmental windows produced lasting changes in male internal and external reproductive anatomy development and hormone levels.

In Experiment 1, when cages were switched to corncob bedding, mice housed on corncob produced significantly fewer pups compared to those housed on cellulose, both at birth and weaning. These findings are consistent with previous reports of endocrine disrupting effects of corncob bedding, including reductions in sexual behavior, disrupted estrous cyclicity, and compromised fertility in rat models (Markaverich et al., 2005; Villalon, Landeros et al., 2012). When all cages were transitioned back to cellulose, reproductive output rebounded among corncob exposed pairs, suggesting that the adverse effects of corncob are at least partially reversible. Pairs previously exposed to corncob outperformed those breeding pairs continuously housed on cellulose, which may seem appealing for boosting breeding productivity, but could actually have detriments to long-term reproductive output. Similar rebound patterns and compensatory effects have been observed following short-term exposure to endocrine disruption or environmental stress and are thought to represent strategies to recover lost reproductive opportunity (Wingfield & Sapolsky, 2003; Metcalfe & Monaghan, 2001). However, this compensatory investment may carry future costs to maternal health or offspring viability, a phenomenon described as “grow now, pay later” (Metcalfe & Monaghan, 2001).

Although the interaction effects in both phases between treatment and litter event were not statistically significant, main effects of both treatment and event were robust and biologically meaningful. Least Significant Number calculations revealed that thousands of litters would be needed to detect any interaction between bedding and litter event, which is unlikely to be practically or biologically relevant. Thus, the primary concern remains the overall reduction in breeding performance associated with corncob bedding, rather than any differences in pup mortality between birth and weaning.

In Experiment 2, we showed that corncob bedding induced changes in male reproductive morphology and endocrine levels that depended on the timing of exposure. Prior work has shown that housing system (IVC vs static cages) interacts with bedding, for example, elevating aggression in IVCs bedded with corncob (Theil et al., 2020). Accordingly, cage ventilation was included as an experimental factor to separate the effects of bedding material from housing system when evaluating endocrine and development disruption in male mice. Among mice housed in IVCs, males had significantly larger AGD when weaned onto corncob bedding than those weaned onto aspen, indicating that IVC housing exacerbated the masculinizing effects during post-weaning corncob exposure. The lack of IVC effect on testosterone *versus* the presence of IVC effect in estrogen and AGD supports aromatase inhibition.

In our study, early-life exposure to corncob was associated with persistently reduced estradiol levels, reduced testosterone levels, reduced AGD, and alterations in baculum morphology, (including increased curvature and reduced shaft extension relative to the bulb). AGD is a well-established marker of sexual development and has been widely used to detect subtle yet biologically meaningful effects of endocrine disruption, with deviations in AGD frequently associated with impaired reproductive function (Schwartz et al., 2019). The reduction in AGD observed following early-life exposure to corncob, with the presence of established endocrine-disrupting compounds in corncob bedding, suggests disruption to normal male development. In contrast, exposure to corncob after weaning (i.e., during sexual maturation) resulted in increased AGD, reflecting a hyper-masculinized phenotype (which aligns with the increase in aggression seen in previous studies). Deviations in AGD reflect disrupted sexual development in males housed on corncob relative to males maintained on aspen throughout development. The direction of this effect depends on the developmental stage of the mouse during exposure.

In addition to changes in AGD, exposure to corncob bedding led to notable changes in baculum morphology. Male mice born on corncob bedding showed greater curvature in both the shaft and bulb of the baculum, as well as a shorter extension of the bulb along the shaft compared to males born on aspen. Altered baculum morphology has direct functional consequences for copulation and reproductive success in rodents, reinforcing the biological relevance of the changes observed in Experiment 2 (Ghione et al., 2025). Importantly, variations in baculum morphology are biologically meaningful, as baculum structure influences copulatory mechanics which contribute to reproductive success in mice under competitive mating conditions (Stockley et al., 2013). In this context, the changes in baculum morphology following early-life corncob exposure may represent an additional pathway through which early endocrine disruption compromises breeding performance.

The endocrine patterns observed in Experiment 2 also provide a potential mechanistic link between corncob bedding exposure and the reduced breeding performance observed in Experiment 1. Testosterone has long been implicated in the regulation of aggressive and reproductive behaviors in mice, particularly through its organizational effects during early development. Importantly, testosterone does not act directly on sex-specific behaviors; rather, its effects depend on conversion to estradiol via aromatase, where activation of estrogen receptor alpha is required for male-typical behaviors in mice (Giammanco et al., 2005). In our study, corncob exposure was associated with reduced circulating testosterone and estradiol, especially during early-life exposure, with the highest hormone levels observed in males housed solely on aspen. Such timing-dependent disruption of hormone signaling during development is likely to have lasting consequences for reproductive behaviors in adulthood.

The clear, timing-dependent effects of corncob bedding on hormone levels, morphology, and breeding performance raise important questions about unrecognized sources of variation in laboratory studies. Our results, in combination with previous findings, show that corncob bedding should be viewed as an uncontrolled extraneous variable in mouse experiments, and should not be used with breeding mice or in studies of reproductive physiology or hormonally sensitive diseases. Furthermore, beyond reproductive and scientific impacts, prior research supports that corncob bedding should be generally avoided for welfare reasons (Mulder, 1975; Krohn & Hansen, 2008; Leys et al., 2012; Theil et al., 2020; Kondo et al., 2022).

### Limitations

Experiment 1 was conducted opportunistically following a sudden change in bedding used within our breeding colony. As a result, breeding pairs were not randomly assigned to bedding groups and animal care personnel were not blinded to treatment conditions. However, due to data mining, the animal care personnel were effectively blinded. Similarly, our data mining was effectively blinded, given the agnostic data processing. Thus, the study is effectively a randomized block design. Our experimental design additionally introduced a potential confound: the reduced breeding performance could be due to the act of changing the type of bedding rather than the biological effects of corncob bedding itself. However, breeding pairs remained on corncob bedding long enough to produce multiple consecutive litters, showing that reduced pup production was not limited to a bedding change. This issue is addressed in Phase 2, once all cages were returned to cellulose bedding. Experiment 1 was performed under real-world institutional conditions, therefore some variation in technician handling and data entry errors may have introduced additional noise. Nonetheless, this variation also enhances the external validity of the findings and their relevance to routine animal breeding operations (Richter et al., 2010).

Although Experiment 2 focused on males, the observed timing-dependent disruption of estradiol signaling raises concern for potential effects on female reproductive function, given the essential role of estrogen and ESR1 signaling in estrous cyclicity, ovarian function, and uterine function (Hamilton et al., 2017). Corncob has widespread effects on physiology and reproduction in females of other species (Markaverich et al., 2005), which warrants future examination in female mice.

### Conclusions

Across two complementary experiments, we found that corncob bedding alters reproductive physiology, affects male sexual development, and impairs breeding performance in laboratory mice. Breeding pairs housed on corncob bedding produced fewer pups than those on cellulose bedding. When returned to cellulose, their breeding performance improved, suggesting a potential compensatory response consistent with prior endocrine disruption. In males of four genetic strains, corncob exposure produced timing-dependent changes in AGD, baculum morphology, and hormone levels, such that early-life exposure reduced masculinization whereas exposure after weaning exaggerated it.

These results speak to the general point that changes in husbandry and operations are all too often driven by engineering solutions in need of a problem, without consideration of the impact on the welfare of the animals involved, or on potential confounds for the science involved. As a whole, these results add to existing literature in other rodents to clearly demonstrate that the use of corncob bedding is fundamentally problematic in terms of animal welfare, physiology, genital malformation, and breeding performance. This conclusion is perhaps best underscored by the fact that corncob is licensed and used as a next-generation rodenticide (Health Canada PMRA, 2025), and that AGD is used in human babies as a measure of exposure to endocrine disruptors with lifelong impacts (Schwartz et al., 2019). As such, we cannot condone the use of corncob as a bedding material for mice (or animals in general).

## Methods

All procedures were approved by Stanford’s Institutional Animal Care and Use Committee. Stanford University is AAALAC accredited.

### Experiment 1

#### Animals

All mice were housed under standard conditions (i.e., 12:12-hour light-dark cycle, at 20-26°C and 30-70% relative humidity). Each breeding cage housed a single male and female in individual ventilated cages (523 cm^2^ floor space; Innovive, San Diego, CA). A total of 420 NSG (NOD SCID gamma) breeding pairs, including multiple NSG substrains, were studied, producing 1,117 litters. Weaning typically occurred around postnatal day 21, with minor variability (up to postnatal day 25). Mice had *ad libitum* access to food (Envigo 2018SX Global Rodent Diet) and water (pre-filled water bottles, Innovive, San Diego, CA). Animals were housed on either corncob bedding (7092-7097 Teklad, Envigo, Madison, WI) or ALPHA-dri® bedding (Shepherd Specialty Papers, Watertown, TN) with nesting material (Enviro-dri, Shepherd Specialty Papers, Watertown, TN). Breeding and husbandry were overseen by Stanford University breeding colony management technicians.

#### Experimental design

Experiment 1 was conducted opportunistically following an unintended change to corncob bedding on a rack within the breeding colony. Mice were housed in two rooms, each containing multiple racks of breeding cages. We opportunistically adopted a quasi-experimental design where one side of each rack was assigned to corncob bedding and the other to cellulose bedding. In other words, while breeding pairs were not individually randomized and personnel were not blinded to treatment, the rack layout resembled a randomized block design. Each rack served as a block, with bedding treatments distributed across sides as evenly and randomly as logistics allowed. In total, 10 rack sides were included in the final analysis.

Data collection ran for 26 weeks and was divided into two 13-week bedding treatment phases of equal duration. Prior to the start of Phase 1, all mice were housed on cellulose bedding. In Phase 1, corncob bedding was introduced to assigned rack sides while other cages remained on cellulose bedding. Given the scale of replacing bedding across two full animal rooms, the transition back to cellulose bedding occurred over several days in the middle of the study and was conservatively categorized as an Intermediate Phase (i.e., bedding was not considered changed for every cage until the end of this phase). Phase 2 then continued with all cages on cellulose bedding for 13 weeks until the study concluded (Fig 1).

**Figure 1.**
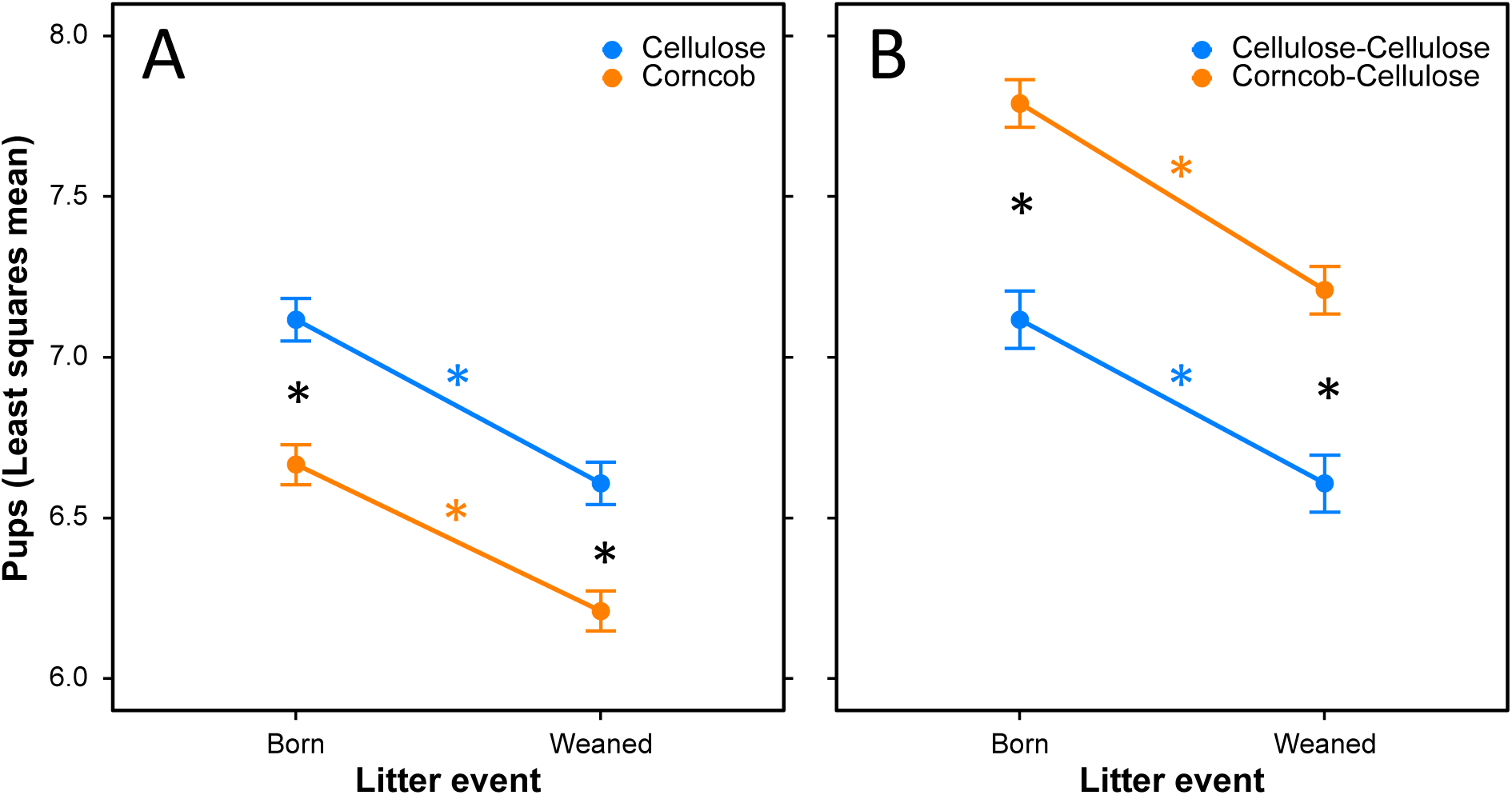
(A) Least Squares Means (± SE) for the number of pups born and weaned per litter for breeding pairs housed on either cellulose or corncob bedding during Phase 1 (N = 324 litters). Significant differences are indicated by *. (B) Least Squares Means (± SE), adjusted for time of year, for number of pups born and weaned per litter during Phase 2 (N = 164 litters).

**Figure 2.**
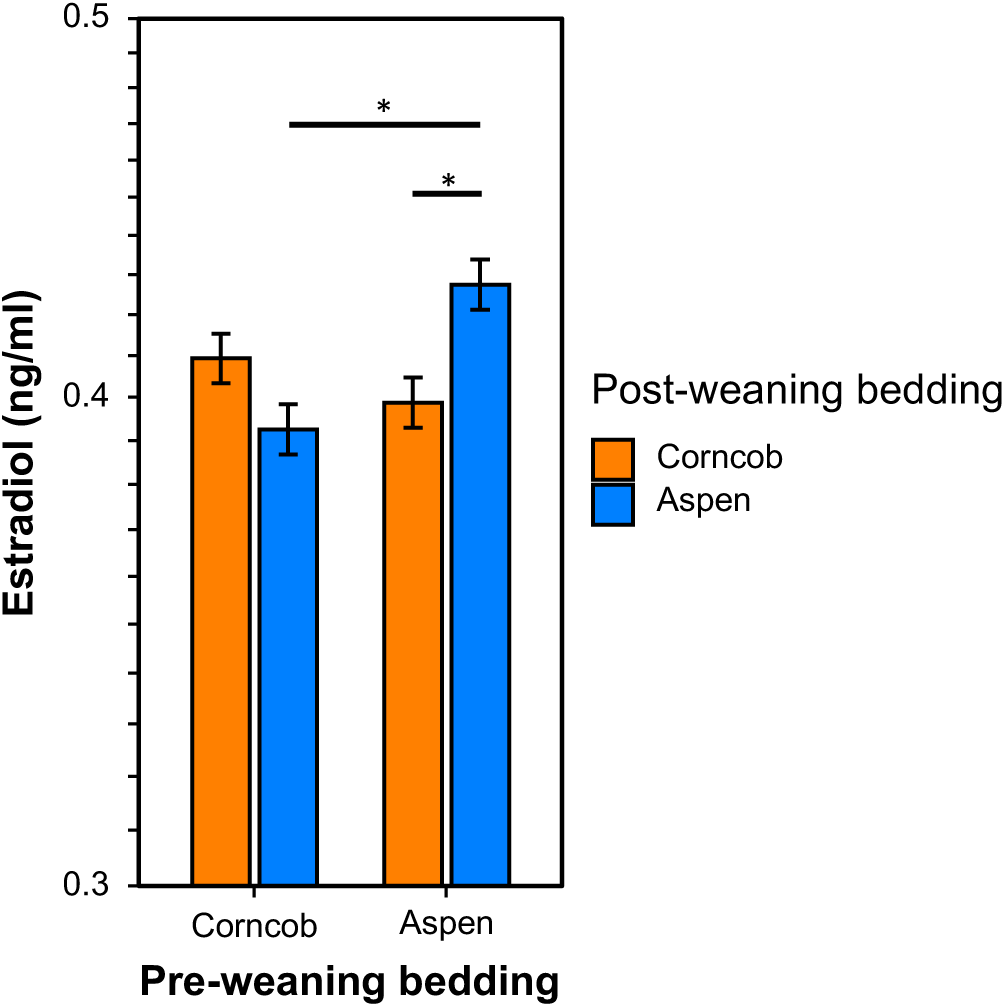
Least Squares Means (± SE) of serum estradiol for males exposed to aspen or corncob bedding before and after weaning (N = 160). Asterisks denote Bonferroni-corrected comparisons (α = 0.025).

**Figure 3.**
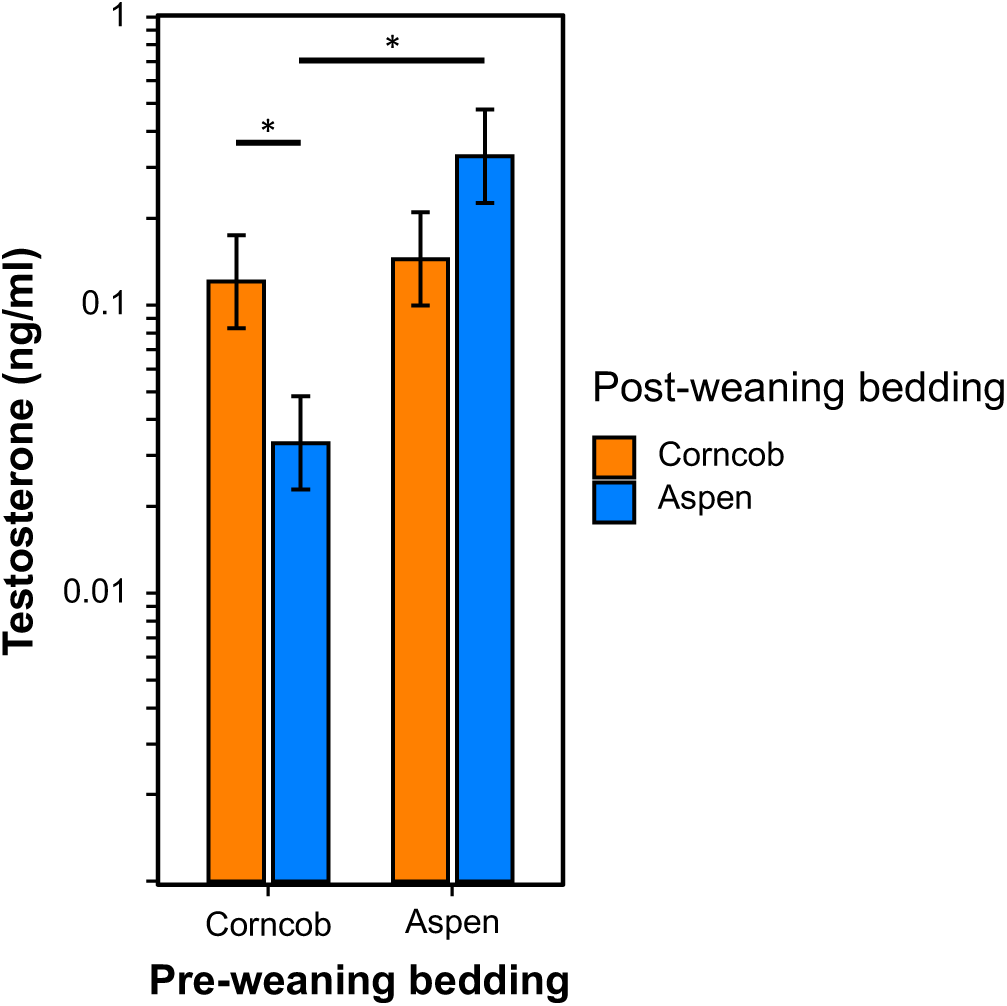
Least Squares Means (± SE) of serum testosterone for males exposed to aspen or corncob bedding before and after weaning (N = 160). Asterisks denote Bonferroni-corrected comparisons (α = 0.025).

Litter sizes and weaning data were recorded by animal care technicians in charge of breeding colony management. Raw breeding records were extracted from the our Colony Management System database, including dates of birth and weaning, pup counts, and associated metadata. Data cleaning was performed using RStudio (version 2024.12.1) with the Tidyverse package (Wickham et al., 2019). Cleaning involved stacking the data, creating breeder- and litter-level summaries, and resolving or excluding litters with missing fields or erroneous data. Litters were treated as the experimental unit with repeated measures on event (birth vs weaning). Thus, litters were excluded from analysis if they had only had one event (i.e., born before the study or weaned after the study), experienced a bedding treatment change between birth and weaning, or were born and weaned in different study phases. Cage cards, technician notes, and electronic records were cross-referenced to correct suspected data entry errors wherever possible; unresolved errors resulted in exclusion of 23 litters. A total of 322 litters were excluded because only one event (birth or weaning) occurred within the study window. After applying all inclusion and exclusion criteria, 488 litters were used in the final analysis.

#### Statistical analysis – Common details and principles

All statistical analyses were conducted in JMP 19, and then replicated in SAS for data deposition (SAS Institute, Cary, NC). All analyses were performed either as General Linear Models (GLM), or as REML Mixed Models. In both cases the assumptions of linear models (homogeneity of variance, linearity, and normality of error) were tested post hoc and suitable transformations applied following best practice (Grafen & Hails, 2002). JMP’s centered polynomial solution was adopted which ensures that marginal terms to interactions represent meaningful tests, non-significant interactions need not be removed, non-significant interactions can be used for data visualization, and post hoc tests of nonsignificant interactions correspond to main effect tests (this last advantage makes data visualization much easier). All data are presented as LSM ± SE.

#### Statistical analysis – Experiment 1

Litter was treated as the statistical unit. Phase 1 and Phase 2 were analyzed independently in order to maximize the number of litters (and breeding pairs) for each comparison. Furthermore, because the Phase 2 analysis focused on breeding pairs that were present in Phase 1 this approach made sure that well-documented seasonal and age-related declines in reproductive performance (e.g., Patel et al., 2017) did not produce spurious results.

For Phase 1, the model included fixed effects for bedding treatment (cellulose or corncob), litter event (birth or weaning), and their interaction. Litter was nested within treatment as a fixed effect to account for repeated measures, allowing each litter to serve as its own control when comparing birth and weaning outcomes. For Phase 2, analysis was limited to breeding pairs present during both phases, comparing those that remained on cellulose (cellulose-cellulose) with those switched from corncob to cellulose (corncob-cellulose). The same general model was used. To account for expected seasonal or age-related effects on decreased reproduction (Patel et al., 2017) in Phase 2, the resulting LSM’s were normalized to the same global baseline, purely for ease of visualization (Figure 1). This post-hoc adjustment does not affect any significance tests.

Bonferroni-corrected post-hoc orthogonal planned contrasts (“slices” in JMP and SAS) were used to examine effects within each litter event and treatment group. Post-hoc power was assessed using the Least Significant Number (LSN) to estimate the number of litters that would be required to detect a significant effect.

### Experiment 2

#### Animals

160 male mice of four strains (C57Bl/6, FVB, CD1, and BALB/c, 40 mice per strain) were used. Mice were housed in single-sex groups of five per cage in individual ventilated cages 523 cm^2^ floor space (Innovive, San Diego, CA) or static cages with static lids, for a total of 32 cages. Given prior data showing corncob effects on aggression in males but not females, this study focused on males (Theil et al., 2020). Mice were housed on either corncob bedding (7092-7097 Teklad, Envigo, Madison, WI) or aspen woodchip (7090 Teklad Sani-Chips, Inotiv, Indianapolis, IN) with nesting material (Enviro-dri, Shepherd Specialty Papers, Watertown, TN) and *ad libitum* access to food (Envigo 2018SX Global Rodent Diet) and water (pre-filled acidified water bottles, Innovive, San Diego, CA). Mice were euthanized at 10 weeks of age using CO_2_. One mouse was euthanized prematurely due to injurious aggression.

#### Experimental design

Sample size for Experiment 2 (N = 32 cages) was determined using Mead’s resource equation (Festing, 2006). We used a 2 x 2 x 2 x 4 factorial design consisting of pre-weaning bedding, post-weaning bedding, cage ventilation, and mouse strain as independent factors (Table 1). Charles River Laboratories bred and housed half of the animals on corncob and half on aspen bedding before shipping to our facility at weaning (approx. 21 days old). Upon arrival, animals were randomly assigned to either remain on their original bedding or were switched to the alternative bedding material, creating four bedding combinations (corncob-corncob, corncob-aspen, aspen-aspen, aspen-corncob). One cage of each strain was assigned to each treatment combination (pre-weaning bedding x post-weaning bedding x ventilation). Half of the animals were housed in static cages and the other half in individual ventilated cages.

**Table 1.** Factorial design of Experiment 2. Four mouse strains were included (C57BL/6, BALB/c, CD1, and FVB). Each group of 4 cages represents one cage per strain. N = 32 cages.

|  | Pre-weaning bedding |  |  |  |
| --- | --- | --- | --- | --- |
|  | Corncob |  | Wood |  |
|  | Static | IVC | Static | IVC |
| Post-weaning bedding |  |  |  |  |
| Corncob | n = 4 cages | n = 4 cages | n = 4 cages | n = 4 cages |
| Wood | n = 4 cages | n = 4 cages | n = 4 cages | n = 4 cages |

#### Baculum imaging

Dissected bacula were scanned using a Bruker SKYSCAN^TM^ 1276 MicroCT at 41.1 μm voxel resolution, with two-frame averaging. Images were reconstructed into axial slices using the SKYSCAN^TM^ software and three-dimension models were generated in RadiANT^TM^. To isolate the baculum, the image window was adjusted per sample until all surrounding tissue artifacts disappeared. Once isolated, each baculum was rotated into the frontal plane for measurement. Six baculum measurements were quantified from each mouse: total baculum length from proximal bulb to distal tip, maximum width of the proximal bulb, minimum width of the shaft, the angle of curvature of the shaft, the angle of curvature of the bulb, and the extension of the bulb along the shaft. The extension of the bulb was defined as the distance from the shaft tip to a line drawn across the widest point of the bulb. For curvature analysis, baculum volumes were exported to DesignSpark Mechanical due to the limitations of RadiANT^TM^ for these measurements. Bacula were oriented into frontal, sagittal, and axial planes. An arc was fitted to the curvature of the shaft in the sagittal plane and to the bulb in the axial plane, and the angle and radius were recorded for each arc.

#### Hormonal assays

Whole blood was collected via cardiac puncture immediately following euthanasia by our necropsy service, and then processed by our diagnostic lab service, following standard protocols for each. Briefly, serum estradiol and serum testosterone concentrations were quantified using commercially available enzyme immunoassay kits run according to kit directions (Arbor Assays DetectX Serum estradiol EIA Kit, Product # KB30; Arbor Assays DetectX Testosterone EIA Kit, Product # K032; Ann Arbor, MI). The estradiol kit had detection limits of 3.75 to 120 pg/ml, with samples diluted 1:20 with assay buffer. The testosterone kit had detection limits of 40.96 pg/ml to 10,000 pg/ml, with samples diluted 1:10 with assay buffer. All samples were assayed in duplicate. Mean intra-assay variation was 7.4% and inter-assay variation across two quality control pools was 9.0%.

#### Statistical analysis

See statistical methods for Experiment 1 for general details. Hormone and AGD data were analyzed using GLM. Serum estradiol and testosterone concentrations were log_10_-transformed to meet the assumptions of linear models. Models included fixed effects of pre-weaning bedding (corncob vs aspen), post-weaning bedding (corncob vs aspen), and their two-way interaction. To account for non-independence among mice housed within the same cage, and the effects of strain, cage was included as a blocking factor nested within the pre-weaning and post-weaning bedding combination. AGD was also log transformed, and the log of bodyweight was included as controlling variable – this allows the model to agnostically identify the correct scaling of AGD for bodyweight (equivalent to a classical allometric log-log regression approach).

To analyze baculum morphology, we adopted a “profile analysis”, an extension of REML mixed models that addresses datasets where the outcome is described by multiple variables (such as morphological measurements, behavioral time budgets, or -omic expression profiles) (Coden et al., 2025). To account for natural differences in baculum shape between mouse strains, all six baculum measurements were standardized as Z-scores within each strain. These standardized measurements were analyzed using a linear mixed model with fixed effects of pre-weaning bedding (corncob vs aspen), dimension (e.g., shaft length, bulb extension, shaft width, bulb width, shaft angle, and bulb angle), and their interaction. Mouse was included as a random effect nested within treatment to account for repeated measures of baculum morphology within individuals. Baculum length (standardized by strain) was included as a covariate to control for any differences in body size. The interaction between Baculum dimension and treatment tests whether the baculum morphology overall is affected by the interacted term. We initially performed the analysis as a fully interactive model. However, this created multiple tests that risk false discovery. Thus, seeing a significant effect of pre-weaning treatment, we reformulated the model as a Hierarchical Linear Model, where all possible treatment effects were first nested within pre-weaning treatment. This approach produces an identical model solution to a fully interactive model, but it explicitly tests whether other treatment combinations offer any additional explanatory value beyond the pre-weaning treatment (the same tests could be performed using post-hoc contrasts on a fully interactive model). This approach tests whether baculum morphology as a whole is affected by treatment in a single test, but further testing of the individual baculum dimensions is not truly hypothesis-driven, and thus these post-hoc tests were Benjamini-Hochberg transformed into q-values (False Discovery Rates) for final interpretation. Effect sizes of each of these comparisons were calculated as standardized mean differences between corncob and aspen, which collapses the dimensionality of the interaction for easier visualization (Figure 5).

**Figure 4.**
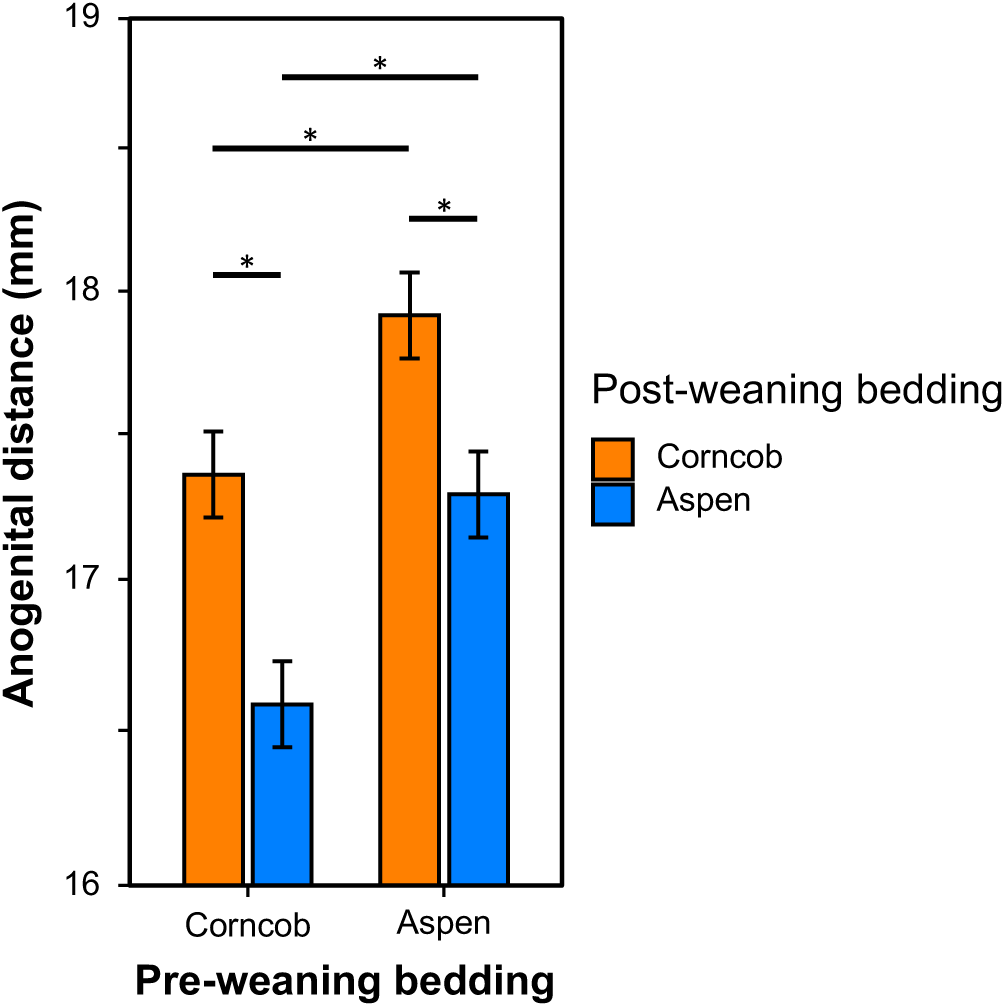
Least Squares Means (± SE) of AGD for males exposed to aspen or corncob bedding before and after weaning (N = 160). Asterisks denote Bonferroni-corrected comparisons (α = 0.025).

**Figure 5.**
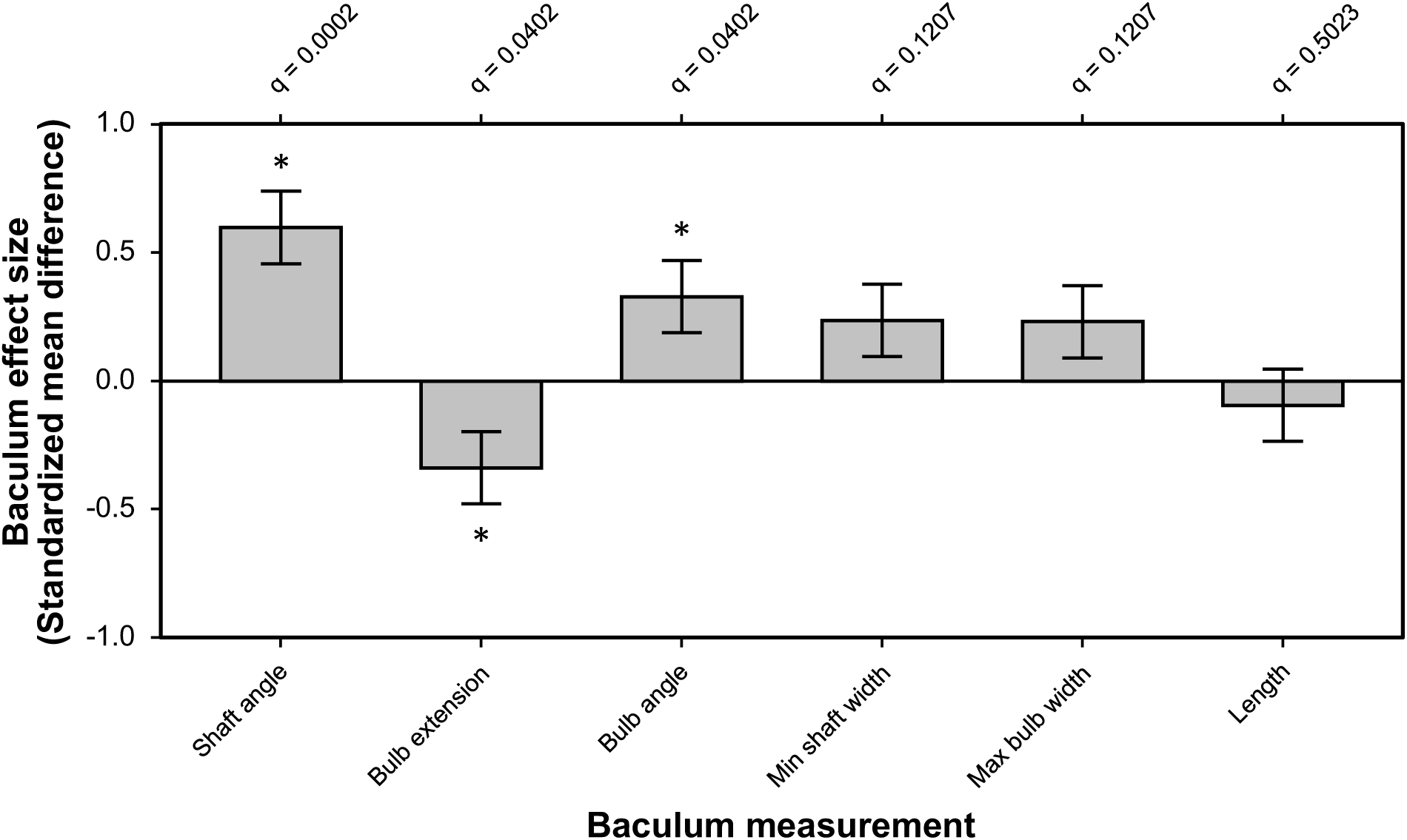
Values represent standardized mean differences (± SE) calculated from z-score measurements within strain. Significant differences are indicated by *. Benjamini-Hochberg corrected q values are plotted on the upper x-axis.

**Figure 6.**
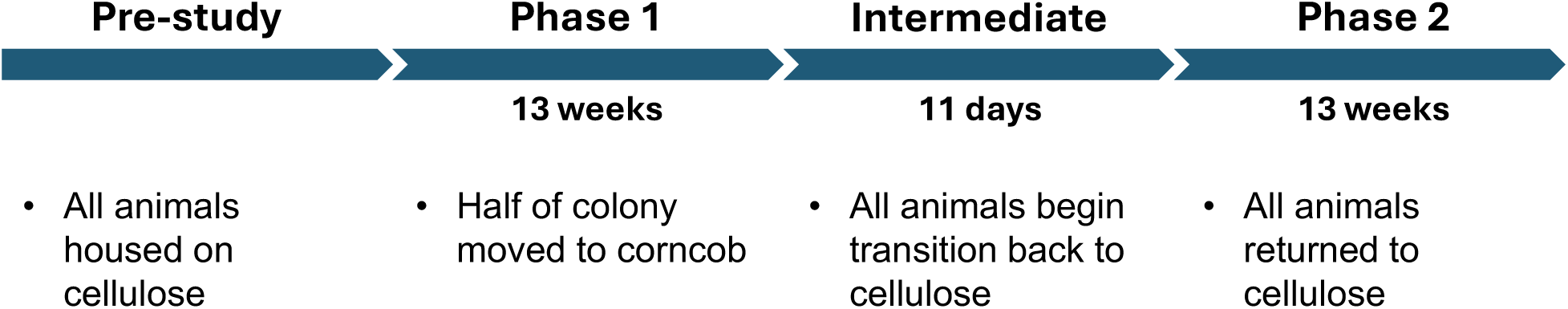
Timeline of bedding exposure across study phases. All breeding cages were housed on cellulose bedding prior to study. During Phase 1, cages were switched to corncob or maintained on cellulose. For Phase 2, all cages returned to cellulose bedding following an intermediate transition period.

## Acknowledgements

We thank the Stanford Veterinary Services breeding colony management staff, Michael Gutierrez, Kaleigh Beacham, Alex Blaney, Jerome Geronimo, and Alexandria Hicks-Nelson for their assistance.

## References

1. Villalon Landeros R, Morisseau C, Yoo HJ, Fu SH, Hammock BD, Trainor BC. 2012. Corncob bedding alters the effects of estrogens on aggressive behavior and reduces estrogen receptor-α expression in the brain. Endocrinology 153(2):949–953. doi:10.1210/en.2011-1745

2. Theil JH, Ahloy-Dallaire J, Weber EM, Gaskill BN, Pritchett-Corning KR, Felt SA, Garner JP. 2020. The epidemiology of fighting in group-housed laboratory mice. Scientific Reports 10(1):16649. doi:10.1038/s41598-020-73620-0

3. Ratuski AS, Theil JH, Ahloy-Dallaire J, Gaskill BN, Pritchett-Corning KR, Felt SA, Garner JP. 2025. Risk factors for barbering in laboratory mice. Scientific Reports 15(1):7456. doi:10.1038/s41598-025-91687-5

4. Kondo SY, Kropik J, Wong MA. 2022. Effect of bedding substrates on blood glucose and body weight in mice. Journal of the American Association for Laboratory Animal Science 61(6):611–614. doi:10.30802/AALAS-JAALAS-22-000047

5. Leys LJ, McGaraughty S, Radek RJ. 2012. Rats housed on corncob bedding show less slow-wave sleep. Journal of the American Association for Laboratory Animal Science 51(6):764–768

6. Mulder JB. 1975. Bedding preferences of pregnant laboratory-reared mice. Behavior Research Methods & Instrumentation 7:21–22. doi:10.3758/BF03201283

7. Krohn TC, Hansen AK. 2008. Evaluation of corncob as bedding for rodents. Scandinavian Journal of Laboratory Animal Science 35:231–236. doi:10.23675/sjlas.v35i4.153

8. Health Canada, Pest Management Regulatory Agency. 2025. Re-evaluation decision RVD2025-09: Cellulose (from powdered corn cobs) and its associated end-use products. Health Canada, Ottawa, Canada.

9. Mani SK, Reyna AM, Alejandro MA, Crowley J, Markaverich BM. 2005. Disruption of male sexual behavior in rats by tetrahydrofurandiols (THF-diols). Steroids 70(11):750–754. doi:10.1016/j.steroids.2005.04.004

10. Markaverich BM, Crowley JR, Alejandro MA, Shoulars K, Casajuna N, Mani S, Sharp J. 2005. Leukotoxin diols from ground corncob bedding disrupt estrous cyclicity in rats and stimulate MCF-7 breast cancer cell proliferation. Environmental Health Perspectives 113(12):1698–1704. doi:10.1289/ehp.8231

11. European Commission. 2024. *Summary report on the statistics on the use of animals for scientific purposes in the Member States of the European Union and Norway in* 2022. Commission Staff Working Document SWD(2024) 185 final. Brussels, Belgium.

12. UK Home Office. 2023. *Annual statistics of scientific procedures on living animals, Great Britain* 2022. Home Office, London, UK.

13. Canadian Council on Animal Care. 2023. *CCAC animal data report* 2022. Canadian Council on Animal Care, Ottawa, Canada.

14. Markaverich B, Mani S, Alejandro MA, Mitchell A, Markaverich D, Brown T, Faith R. 2002. A novel endocrine-disrupting agent in corn with mitogenic activity in human breast and prostatic cancer cells. Environmental Health Perspectives 110(2):169–177. doi:10.1289/ehp.02110169

15. Baskin L, Sinclair A, Derpinghaus A, Cao M, Li Y, Overland M, Cunha GR. 2021. Estrogens and development of the mouse and human external genitalia. Differentiation 118:82–106. doi:10.1016/j.diff.2020.09.004

16. Ryan BC, Vandenbergh JG. 2002. Intrauterine position effects. Neuroscience & Biobehavioral Reviews 26:665–678. doi:10.1016/S0149-7634(02)00038-6

17. Schwartz CL, Christiansen S, Vinggaard AM, Axelstad M, Hass U, Svingen T. 2019. Anogenital distance as a toxicological or clinical marker for fetal androgen action and risk for reproductive disorders. Archives of Toxicology 93(2):253–272. doi:10.1007/s00204-018-2350-5

18. Ghione CR, Schultz NG, Park S, Menke DB, Dean MD. 2025. Genetic disruption of the baculum compromises the ability of male mice to copulate. PLoS Genetics 21(7):e1011787. doi:10.1371/journal.pgen.1011787

19. Stockley P, Ramm SA, Sherborne AL, Thom MD, Paterson S, Hurst JL. 2013. Baculum morphology predicts reproductive success of male house mice under sexual selection. BMC Biology 11:66. doi:10.1186/1741-7007-11-66

20. Wingfield JC, Sapolsky RM. 2003. Reproduction and resistance to stress: when and how. Journal of Neuroendocrinology 15(8):711–724. doi:10.1046/j.1365-2826.2003.01033.x

21. Metcalfe NB, Monaghan P. 2001. Compensation for a bad start: grow now, pay later? Trends in Ecology & Evolution 16(5):254–260. doi:10.1016/S0169-5347(01)02124-3

22. Richter SH, Garner JP, Auer C, Kunert J, Würbel H. 2010. Systematic variation improves reproducibility of animal experiments. Nature Methods 7(3):167–168. doi:10.1038/nmeth0310-167

23. Giammanco M, Tabacchi G, Giammanco S, Di Majo D, La Guardia M. 2005. Testosterone and aggressiveness. Medical Science Monitor 11:RA136–RA145.

24. Wickham H, Averick M, Bryan J, Chang W, McGowan L, François R, Grolemund G, Hayes A, Henry L, Hester J, Kuhn M, Pedersen T, Miller E, Bache S, Müller K, Ooms J, Robinson D, Seidel D, Spinu V, Takahashi K, Vaughan D, Wilke C, Woo K, Yutani H. 2019. Welcome to the tidyverse. Journal of Open Source Software 4:1686. doi:10.21105/joss.01686

25. Grafen A, Hails R. 2002. Modern statistics for the life sciences. Oxford University Press, Oxford, UK.

26. Patel R, Moffatt JD, Mourmoura E, Demaison L, Seed PT, Poston L, Tribe RM. 2017. Effect of reproductive ageing on pregnant mouse uterus and cervix. Journal of Physiology 595(6):2065–2084. doi:10.1113/JP273350

27. Festing MF. 2006. Design and statistical methods in studies using animal models of development. ILAR Journal 47(1):5–14. doi:10.1093/ilar.47.1.5

28. Hamilton KJ, Hewitt SC, Arao Y, Korach KS. 2017. Estrogen hormone biology. In: Forrest D, Tsai S, editors. Current Topics in Developmental Biology. Vol 125. Academic Press. p.109–146.

29. Coden KM, Beacham KJ, Stix-Brunell BE, Moorhead R, Byrd KA, Baker JN, Garner JP. 2025. Stereotypy is strongly linked to multiple biomarkers of oxidative stress—A potential common etiology for abnormal repetitive behaviors. PLOS ONE 20(11):e0326902. doi:10.1371/journal.pone.0326902

30. Allen PS, Lawrence J, Stasula U, Pallas BD, Freeman ZT. 2021. Effects of compressed paper bedding on mouse breeding performance and recognition of animal health concerns. Journal of the American Association for Laboratory Animal Science 60(1):28–36. doi:10.30802/AALAS-JAALAS-20-000036

31. Bratcher NA, Allen CM, McLahan CL, O’Connell DM, Burr HN, Keen JN, Burns MA. 2022. Identification of rodent husbandry refinement opportunities through benchmarking and collaboration. Journal of the American Association for Laboratory Animal Science 61(6):624–633. doi:10.30802/AALAS-JAALAS-21-000099

32. Burn CC, Mason GJ. 2005. Absorbencies of six different rodent beddings: commercially advertised absorbencies are potentially misleading. Laboratory Animals 39(1):68–74. doi:10.1258/0023677052886592

33. Domer DA, Erickson RL, Petty JM, Bergdall VK, Hickman-Davis JM. 2012. Processing and treatment of corncob bedding affects cage-change frequency for C57BL/6 mice. Journal of the American Association for Laboratory Animal Science 51(2):162–169

34. Hashikawa K, Hashikawa Y, Tremblay R, Zhang J, Feng JE, Sabol A, Lin D. 2017. Esr1⁺ cells in the ventromedial hypothalamus control female aggression. Nature Neuroscience 20(11):1580–1590. doi:10.1038/nn.4644

35. Jones SL, Antonie RA, Pfaus JG. 2015. The inhibitory effects of corncob bedding on sexual behavior in the ovariectomized Long-Evans rat treated with estradiol benzoate are overcome by male cues. Hormones and Behavior 72:39–48. doi:10.1016/j.yhbeh.2015.05.002

36. Matsumoto T, Honda S, Harada N. 2003. Alteration in sex-specific behaviors in male mice lacking the aromatase gene. Neuroendocrinology 77(6):416–424. doi:10.1159/000071313

37. Markaverich BM, Alejandro M, Thompson T, Mani S, Reyna A, Portillo W, Crowley JR. 2007. Tetrahydrofurandiols (THF-diols), leukotoxindiols (LTX-diols), and endocrine disruption in rats. Environmental Health Perspectives 115(5):702–708. doi:10.1289/ehp.9311

38. Markaverich BM, Alejandro MA, Markaverich D, Zitzow L, Casajuna N, Camarao N, Crowley JR. 2002. Identification of an endocrine disrupting agent from corn with mitogenic activity. Biochemical and Biophysical Research Communications 291(3):692–700. doi:10.1006/bbrc.2002.6499

39. Tataryn NM, Buckmaster CA, Schwiebert RS, Swennes AG. 2021. Comparison of four beddings for ammonia control in individually ventilated mouse cages. Journal of the American Association for Laboratory Animal Science 60(1):37–43. doi:10.30802/AALAS-JAALAS-20-000051

40. Vandenberg LN, Colborn T, Hayes TB, Heindel JJ, Jacobs DR, Lee DH, Myers JP. 2012. Hormones and endocrine-disrupting chemicals: low-dose effects and nonmonotonic dose responses. Endocrine Reviews 33(3):378–455. doi:10.1210/er.2011-1050

41. Wu MV, Manoli DS, Fraser EJ, Coats JK, Tollkuhn J, Honda S, Shah NM. 2009. Estrogen masculinizes neural pathways and sex-specific behaviors. Cell 139(1):61–72. doi:10.1016/j.cell.2009.07.036

42. Wu MV, Tollkuhn J. 2017. Estrogen receptor alpha is required in GABAergic, but not glutamatergic, neurons to masculinize behavior. Hormones and Behavior 95:3–12. doi:10.1016/j.yhbeh.2017.07.001

